# Modelling inflammation-induced peripheral sensitization in a dish – more complex than expected?

**DOI:** 10.1101/2024.08.12.607558

**Authors:** Yuening Li, Amy Lock, Laura Fedele, Irene Zebochin, Alba Sabate, Matthew Siddle, Silvia Cainarca, Pascal Röderer, Katharina Montag, Paola Tarroni, Oliver Brüstle, Tanya Shaw, Leonie Taams, Franziska Denk

**Author notes:** These authors have contributed equally to this work (Yuening Li, Amy Lock, Laura Fedele).

## Abstract

Peripheral sensitization of nociceptors is believed to be a key driver of chronic pain states. Here, we sought to study the effects of a modified version of inflammatory soup on the excitability of human stem-cell derived sensory neurons. For this, we used a pre-existing and a novel stem cell line, modified to stably express the calcium sensor GCamP6f.

Upon treatment with inflammatory soup, we observed no changes in neuronal transcription or functional responses upon calcium imaging, and only a very minor increase in resting membrane potential via whole cell patch clamping. Similarly small changes were observed when treating mouse primary sensory neurons with inflammatory soup. A semi-systematic re-examination of past literature further indicated that observed effects of inflammatory mediators on dissociated sensory neuron cultures are generally very small.

We conclude that modelling inflammation-induced peripheral sensitization *in vitro* is non-trivial and will require careful selection of mediators and/or more complex, longitudinal multi-cellular setups. Especially in the latter, our novel GCamP6f induced-pluripotent stem cell line may be of value.

## Introduction

Peripheral sensitization is the process by which sensory neurons alter their firing properties upon inflammation or injury, becoming more easily triggered in response to their usual ligands and activators [9; 25]. The latter include hot or cold temperatures, noxious mechanical activation, low pH, damaging chemicals like capsaicin and endogenously produced pro-inflammatory molecules, like cytokines and growth factors. Given the importance of peripheral sensitization in a wide variety of painful conditions, including neuropathies [17], arthritis [35; 45] and immune-mediated inflammatory diseases [15], it is not surprising that many have tried to model it in an *in vitro* setting.

Early seminal studies by Handwerker, Reeh and their teams used rodent skin-nerve explants to study the effects of what they called ‘inflammatory soup’ (IS) – four putative pro-algesic molecules known to be present in an inflammatory environment: histamine, bradykinin, serotonin (5-HT) and prostaglandin-E2 (PGE2). They were able to demonstrate increased nociceptive neuron firing in the presence of IS, which was significantly potentiated by low pH [41]. They also went on to show that IS causes pain when perfused through the skin of healthy volunteers [40].

Since then, IS and its components have also been studied in dissociated dorsal root (DRG) and trigeminal ganglion (TG) cultures, where they are reported to alter neuronal hyperexcitability, e.g. [5; 21; 30; 38; 46]. Most of the work has been conducted in rodent cells with a few notable exceptions [11; 47].

Here we set out to study the effects of IS in human induced pluripotent stem cell (iPSC)-derived sensory neurons (iSNs). We used the Chambers protocol [7] for differentiation with small molecules and a modified version of the original IS, adding tumor necrosis factor alpha (TNF) and nerve growth factor (NGF), in addition to 5-HT, bradykinin, histamine and PGE2. This was to ensure that our inflammatory solution would contain no shortage of molecules which can bind iSNs directly. For example, RNA sequencing (RNA-seq) suggests that bradykinin receptors are only expressed at very low levels in human post-mortem DRG [18] and absent from our iSNs (see Results). Similarly, PGE2 can bind four different receptors, two of which are clearly expressed in postmortem DRG (PTGER3 and PTGER4). They are also the two which have been claimed to be most important for sensory neuron sensitization [39]. In contrast, iSNs express only PTGER4 at rather low levels. Interestingly, the functional data on these two molecules is inverse to their neuronal receptor expression: bradykinin has consistently been shown to be painful when infused into skin [31; 40], while PGE2 reportedly does not induce pain beyond that associated with fluid injection [8], unless potentiated by low pH [33].

We assessed the consequences of modified-IS on our human sensory neurons using RNA-seq, calcium imaging and patch clamping. We found its effects to be surprisingly modest and propose that this might be a general issue: a re-examination of prior literature, as well as our own experiments in mouse DRG neuron cultures indicate that studying peripheral sensitization in dissociated cells may not be as straightforward as conventionally assumed.

## Methods

### Generation of GCaMP6f iPSC line

The iPSC-line UKBi013-A (https://hpscreg.eu/cell-line/UKBi013-A) was generated via Sendai virus reprogramming of peripheral blood mononuclear cells, obtained from a healthy male donor. The use and generation of iPSC lines was approved by the Ethics Committee of the Medical Faculty of the University of Bonn (approval number 275/08), and informed consent was obtained from the donor. The human UKBi013-A-GCamP6f iPSC reporter line for functional testing was generated by genome editing using CRISPR/Cas9 technology. The region to be modified, here AAVS1, was PCR-amplified from a genomic DNA sample of UKBi013-A and sequenced. Variations from the NCBI reference genome sequence were recorded and incorporated into the targeting vectors. The following constructs were designed and generated: (1) a Cas9 expression construct driven by the elongation factor (EF) 1α promoter, (2) single guide RNA-expressing constructs specific for the targeted locus under control of a U6 RNA polymerase promoter, and (3) a homology-directed repair (HDR) donor plasmid, AAVS1-GCaMP6f, containing the insertion cassette flanked by locus-specific homology arms. All constructs were verified by sequencing and positively tested in proof-of-concept studies in HEK293 cells. The human iPSC line UKBi13A was cultured and maintained in StemMACS iPS-Brew (Miltenyi Biotec, #130-104-368) and on Geltrex (Thermo Fisher Scientific, #A1413301) coated plates. Cells were passaged every 3-4 days using 0.5mM EDTA. Single cell suspensions for transfection experiments were obtained after enzymatic treatment with Accutase (Thermo Fisher Scientific, #A1110501). Nucleofections were performed with a 4D-Nucleofector System (LONZA) using the Primary P3 kit according to the manufacturer’s instructions (program used: CB-150). The following plasmids were co-transfected: the Cas9 expression construct, an AAVS1-targeting sgRNA expression construct, and the AAVS1-GCaMP6f HDR donor vector. After 48 hours of cell recovery, puromycin was applied to select for positive clones. After selection, individual clones were picked manually, expanded and screened by PCR for positive modification of the AAVS1 locus. For positive clones, PCR products were sequenced and analysed to verify the correct editing of the targeted locus.

### Human iPSC-derived sensory neuron differentiation

Experiments were conducted with two iPSC lines derived from one healthy male and one healthy female human donor: HPSI0714i-kute_4 (Kute4), female origin, obtained via the Human Induced Pluripotent Stem Cell (HIPSCI) Initiative at King’s College London and UKBi013-A-GCamP6f (UKB), male origin. iPSCs were maintained on 6-well plates coated with vitronectin (StemCell Technologies, catalogue #100-0763) in Stemflex media (ThermoFisher, #A3349401) until 80% confluence. Cells were gently dissociated from wells using Versene (ThermoFisher, #15040066) for 4 mins at 37°C. Prior to differentiation, iPSCs were seeded on 6-well plates coated with Geltrex (ThermoFisher, #A1413302) and allowed to expand in Stemflex media to 60% confluence. At this point (day 1), the medium was changed to mouse embryonic fibroblast conditioned medium (MEF, Bio-Techne, #AR005) supplemented with 10ng/mL human recombinant FGF-2 (Miltenyi, #130-093-839). 24 hours later (day (D)0), differentiation was initiated according to the Chambers protocol [7]. Briefly, from D0-D3, KSR media (Knockout DMEM (ThermoFisher, #10829018), 15% knockout serum replacement (ThermoFisher, #10828010), 1% Glutamax (ThermoFisher, #35050038), 1% non-essential amino acids (ThermoFisher, #11140035), 100uM beta-mercaptoethanol (ThermoFisher, #31350010), 1% antibiotic/antimycotic (ThermoFisher, #15240096) was added and changed daily. From D0-D1, SMAD inhibitors (10µM SB431542, Bio-techne, #1614-10; 100nM LDN 193189, Bio-techne, #6053-10) were added to media. From D2-D3, in addition to SMAD inhibitors, 3µM CHIR99021 (Bio-techne, catalogue #4423-10), 10µM SU5402 (Merck, #SML0443), 10µM DAPT (Bio-techne, #2634-10) were also added to media. From D4-D5, media composition was changed to 75% KSR media + 25% N2 media (Neurobasal medium (ThermoFisher, #21103049), 2% B27 supplement (ThermoFisher, #17504044), 1% N2 supplement (ThermoFisher, #17502048), 1% Glutamax, 1% antibiotic/antimycotic) with all 5 inhibitors included. From D6-7, media composition was changed to 50% KSR + 50% N2, and only 3µM CHIR99021, 10µM SU5402 and 10µM DAPT included. From D8-9, media composition was changed to 25% KSR + 75% N2, and only 3µM CHIR99021, 10µM SU5402 and 10µM DAPT included. On D10, media composition was changed to 100% N2, and only 3µM CHIR99021, 10µM SU5402 and 10µM DAPT included. On D11, immature neurons were dissociated from wells using TrypLE (ThermoFisher, #12604013) for 7 mins at 37°C. Neurons were replated to Geltrex-coated 13mm glass coverslips at a density of 75,000 cells/well. Neurons were matured and maintained in N2 media containing 25ng/mL human β-NGF (Peprotech, #450-01), 25ng/mL human NT3 (Peprotech, #450-03), 25ng/mL human GDNF (Peprotech, #450-10), 25ng/mL human BDNF (Peprotech, #450-02). From D11-D14, media also contained 10μM ROCKi (Enzo, #ALX-270-333-M005) and 3µM CHIR99021. Media was changed twice a week. To remove proliferating, non-neuronal cells, 1-2µM Cytosine beta-D-arabinofuranoside (AraC, SLS, #C1768-100MG) was added every other media change until non-neuronal cells were removed. Neurons were matured for different lengths depending on each experimental question – see other method sections and figure legends.

### Mouse dorsal root ganglion (DRG) cultures

All animal work was conducted in accordance with UK Home Office Legislation (Scientific Procedures Act 1986) and approved by the Home Office to be carried out at King’s College London under a current project license. Cultures of dissociated dorsal root ganglia (DRG) were prepared from male and female adult C56BL/6J mice purchased from Charles River. Mice were terminally anaesthetised with pentobarbital and transcardially perfused with 10ml of ice-cold Dulbecco’s phosphate-buffered saline (DPBS). Within 1 hour, as many DRG as possible were collected from cervical, thoracic and lumbar levels. They were placed into dissociation buffer made up of F12 media (Sigma-Aldrich, #N6658) containing 2.5 mg/ml collagenase (Sigma Aldrich, C9722) and 5 U/ml Dispase II (Sigma Aldrich, #D4693). After a 45-minute digestion at 37°C in a 5% CO_2_ incubator, DRG were triturated with a P1000 pipette and passed through a cell strainer. They were then resuspended in DRG media (F12 with 1% antibiotic/antimycotic (Thermofisher, #15240096), 1% N2 supplement (ThermoFisher, #17502048), 10% FBS (Gibco, #10500-064) and plated at a density of 5,000 cells per 24-well on poly-L-lysine (Sigma-Aldrich, #P8954) and Matrigel (Scientific laboratory supplies, #356230) coated coverslips. Patch clamp recordings were performed after 3-7 days in culture (see figure legends for more detail).

### Immunostaining

Coverslips containing sensory neurons (Day 46-81) were fixed in 2% paraformaldehyde (PFA) for 10 minutes at 37⁰C. They were rinsed twice with PBS and kept in PBS until staining. Coverslips were blocked for 1h in PBS-Triton (0.2% Triton X-100, Sigma-Aldrich, #T-9284) with 10% Normal Donkey Serum (NDS, Abcam, #AB7475) at room temperature. They were then incubated at room temperature overnight with primary antibodies: 1:100 Brn3a (Sigma-Aldrich, #MAB1585) 1:500 PGP9.5 (Abcam, #ab108986), 1:300 NeuN (Cell Signalling, #12943S) in PBS-Triton + 10% NDS.

On the second day, the coverslips were washed 3x with PBS prior to incubation with secondary antibodies (1:1000 donkey anti-mouse AF488 Invitrogen, 1:1000 donkey anti-rabbit AF568 Invitrogen) in PBS-Triton + 10% NDS for 2h at room temperature. They were then washed 3x in PBS and mounted on glass slides using DAPI-containing mounting medium (DAPI Fluoromount-G, #0100-20). The slides were left to dry overnight at room temperature prior to imaging.

### Imaging and analysis

The slides were imaged using a Zeiss LSM 710 confocal microscope with a 20x objective. A minimum of three coverslips per trial were imaged. One representative picture per coverslip was analysed using FIJI [36], using a novel macro, which we have made freely available on Zenodo, alongside more detailed instructions (https://doi.org/10.5281/zenodo.12783430). Briefly, we first obtained a maximum intensity projection from the raw image. The channels were then split, and a binary mask of each channel was obtained using the “Threshold” function. The “Analyze Particle” function was used to calculate the total number of DAPI+ nuclei. All the channels were then analyzed in pairs using the “Colocalization Threshold” function followed by “Colour Threshold”. Since all the antibodies we used stained the nucleus (either exclusively or not), the resulting “regions of interest” (ROIs) represented the total number of co-localized nuclei detected. To obtain the final number of double positive nuclei, we excluded any ROI with area <30µm to avoid any dead cells and background specks. Occasionally two or more nuclei were counted by the programme as a single ROI, hence we considered areas above 130 µm to be two nuclei and above 200µm to be three joined nuclei.

### Calcium imaging and analyses

Neurons derived from Kute4 or UKB lines over 50 days old were considered functionally mature and used in calcium imaging. Kute4 neurons were pre-incubated with 2.5 μg/μl Fura2-AM and 1uM probenecid (Sigma P8761) for 1-1.5 hours in an incubator (37°C, 5% CO_2_). For GCaMP6f+ UKB neurons, coverslips were habituated in extracellular solution (ECS) for about 1h before usage. During the experiment, the coverslip was perfused with ECS containing: 140 mM NaCl (VWR, #7647-14-5), 5 mM KCl (Fisher Scientific, #7447-40-7), 10 mM glucose (Sigma, #G7021), 10 mM HEPES (Fisher Scientific, #BP310-500), 2 mM CaCl_2_ (Sigma, #C-7902) and 1 mM MgCl_2_ (BDH, #290964Y). The pH was adjusted to 7.4 with 1M NaOH. High KCl solution contained 95 mM NaCl, 50 mM KCl with the other ingredients identical, pH adjusted to 7.4. The perfusion speed was set to be around 4 mL/min. Pregnenolone sulphate (Sigma, #P162) and veratridine (ENZO, #BML-NA125-0010) were reconstituted according to the manufacturers’ instruction and diluted in ECS to use at the final concentration indicated in the figures.

Calcium imaging was performed at room temperature on an inverted microscope (Nikon Eclipse TE200 microscope), equipped with a 10x air objective (NA 0.8) and a high-speed random-access monochromator (Photon Technology International) for the light source, and an ORCA-flash4.0 camera. To record Fura2 signal, measurements of 340 nm, 380 nm and their ratio (340 nm/380 nm) were obtained by the Easyratio2 software. To record GCaMP6f signalling, measurements at 488 nm were obtained.

The time-lapse data were exported as videos in TIF files and then analysed in FIJI. To segment and select individual neurons, the StarDist plug-in [37] was used to identify individual cells as ROIs followed by manual inspection. The measurement values at different frames were exported using the multi-measure function. Custom-made R code, made freely available on Zenodo (https://doi.org/10.5281/zenodo.13122856), was used to extract the mean value for each frame for each ROI. Specifically, ΔF/F was calculated by subtracting the reading for each time frame with the baseline average and divided by the baseline reading. A second baseline was calculated in the wash time just before 50 mM KCl. Live cells were defined from the ΔF/F value during the KCl period: at least 10 frames had to exceed the second baseline by at least 3 standard deviations (SD). Within the live-cell subset, PS and veratridine responders were defined as cells in which the ΔF/F values exceeded the baseline mean by at least 3 SD for a minimum of 80 data frames (∼2 minutes). The response also needed to occur within 5 min of the compound being added and be at least 10% above the baseline for a cell to be counted as a responder. The percentage of live cells and of drug responders was calculated. The max % response was obtained by averaging the 3 maximal values of drug responders over the response timeframe. Plots were created in R and Prism 9 GraphPad software.

### Whole-cell patch clamp electrophysiology

Both iPSC-derived neurons and neurons dissociated from mouse DRG were continuously superfused with bicarbonate buffered solution containing in mM: 124 NaCl (Acros Organics, #207790250), 2.5 KCl (Thermoscientific, #196770010), 26 NaHCO_3_ (Sigma-Aldrich, #71631), 1 NaH_2_PO_4_ (Sigma-Aldrich, #71500), 2 CaCl_2_ (Honeywell Fluka, #21114), 1 MgCl_2_ (Honeywelll Fluka, #63020), 10 glucose (Sigma-Aldrich, #G8270), pH 7.4, bubbled with 95% O_2_ and 5% CO_2_. Patch pipettes solution consisted of (mM): 130 KGluconate (Acros Organics, #229322500), 2 NaCl (Acros Organics, #207790250), 0.01 CaCl_2_ (Honeywell Fluka, #21114), 2 MgCl_2_ (Honeywelll Fluka, #63020), 10 HEPES (Sigma-Aldrich, #H3375), 0.1 EGTA (Bio Basic, #ED0077), 2 ATP-Na (Sigma-Aldrich, #A7699), 0.5 GTP-Na (Sigma-Aldrich, #G8877), 10 Phospocreatine-Na (Sigma-Aldrich, #P7936), osmolarity 280 ± 10, adjusted to pH 7.2 with 1M KOH (BDH, #29627). The liquid junction potential was -14mV and adjusted offline. DRG (3-7 days in culture) or iSNs (Day 52-63 Kute4; Day 45-55 UKB) were employed for electrophysiological experiments. To avoid any confounding effect of freshly added growth factors, media of iSNs were not changed for 3 days before the experiments. Coverslips were incubated for 18-26h with either control media (Ctrl: DRG media without FBS, N2 media without growth factors) or modified inflammatory soup (modified-IS) media (Ctrl media with 100 ng/ml NGF (Peprotech, #450-01-100), 100 ng/ml TNF (Peprotech, #300-01A-100), 1 μM histamine (Sigma-Aldrich, #H7250), 1 μM bradykinin (Sigma-Aldrich, #B3259), 1 μM serotonin (Sigma-Aldrich, #H9523), 1 μM prostaglandin E2 (Bio-techne, #2296). All electrophysiological experiments were undertaken at room temperature. Patch pipettes were used at a resistance of 4.5-6.5 MΩ and were pulled using filamented borosilicate glass (HARVARD GC150F-10, #30-0057), with an outer and inner diameter of 1.5 mm and 0.86 mm, respectively. Whole-cell patch current clamp recordings were undertaken using a MultiClamp 700B amplifier (Molecular Devices), digitised using a Digidata 1440A (Molecular Devices), with an upright microscope Olympus BX51WI. Recordings were acquired and analysed with the pClamp software suite v.11.3. Five iSN differentiations batches were used with a total of 45 cells in each treatment group. Six mouse DRG cultures (3 males and 3 females) were employed with a total of 26-29 cells in each treatment group. Spontaneous action potentials were recorded in gap-free mode (2 minutes) at resting membrane potential. A cell was defined as firing spontaneous action potentials if it fired at least one action potential during this period. Evoked action potentials were recorded at a 0.5 Hz frequency using a 200ms current injection from -50 pA up to 850 pA with a stepwise 25 pA increase, both at resting membrane potential and after injecting currents to hold the cells -75 mV. Cells that did not evoke an action potential upon current injection (up to 850 pA) or with a resting membrane potential above -40 mV were excluded from our final analysis. Resting membrane potential was calculated as an average of the first 200 ms before the stimulus protocol. Rheobase was measured at resting membrane potential and was defined as the minimum injected current that would elicit an action potential. Action potential amplitude, threshold, half-width and afterhyperpolarisation (fast and medium) were measured holding cells at -75 mV. The peak amplitude was measured between the resting membrane potential (-75 mV) and the peak. Action potential threshold was defined using the first derivative method, when the membrane potential crossed dV/dt equal 10 mV/ms [32]. Half-width is the duration of the action potential at half of its peak amplitude. Fast and medium afterhyperpolarization are respectively the minimum potential following the first action potential peak and the minimum potential following the stimulus protocol. Time to peak was measured as the time between the stimulus and the first action potential peak. To assess the firing frequency (number of action potentials) upon current injection, an action potential was defined as such if dV/dt was equal or higher than 10 mV/ms.

### RNA sequencing

Two batch-controlled sequencing experiments were conducted, comparing iSNs at three different timepoints (day 29-30, day 50-53, day 69-70) and comparing day 60 neurons treated with modified-IS versus control media for 24 hours. For each group, two coverslips from a 24-well plate of neurons were combined into 350 µL RLT buffer from the RNeasy Micro Plus kit (Qiagen, #74134), supplemented with 1% beta-mercaptoethanol (PanReac AppliChem, #A1108.0100). The samples were stored at -80 °C degrees, until batch-controlled RNA extraction, library preparation and sequencing. RNA was purified using the RNeasy Micro Plus kit (Qiagen, #74134) following manufacturer’s instructions. We quantified the RNA amount using a Qubit high sensitivity RNA assay (Invitrogen, #Q32851) and checked its quality using a Tapestation 4150 system (Agilent). All samples had a RIN value of more than 9.9. They were processed at once within the same library preparation and multiplexed into the same Illumina sequencing lane (150bp, paired-end reads) by the service provider Novogene Sequencing.

Reads were pseudo-aligned with kallisto version 0.50.1 to the human genome: Homo Sapiens GCRh38, kallisto index version 13 [3]. An average of 32M reads were sequenced per sample, of which an average of 28M reads were successfully aligned. See **Supplementary Table 1** for alignment statistics for each sample. Genes were considered to be expressed in our dataset if their Transcripts Per Million (TPM) value was equal to 1 or more in all samples of a given group. Differential expression was performed by running DESeq2 [24] in R. All fastq and processed files are available on the Gene Expression Omnibus (GEO) repository under accession number GSE268585.

Pseudobulk count data were obtained from previously published single-nucleus RNA-seq data [18; 29] using GEO accession numbers GSE168243 & GSE201586. The Jung et al dataset [18] was subset on their DRG neuron cluster, while the Nguyen et al [29] GEO deposition only contained the neuronal fraction of their sequencing data. In both cases, counts were normalised in Seurat v0.4 [14] using the ‘RC’ method and summed across all nuclei within a sample. Here, we display data for all n = 6 separate samples in Nguyen et al. and n = 16 out of 18 samples in Jung et al., since two of the latter (original IDs: SAM24364375, SAM24364376) only contained sequencing data on 20 vs. 6 nuclei each and thus were deemed too unreliable for pseudo-bulk generation. Please note that both datasets are somewhat contaminated with glial cell transcripts, like SOX10, COL15A1 and FABP7 detectable at varying levels. FABP7 ranked 6898^th^ in Nguyen et al. & 233^th^ in Jung et al. in terms of average expression across samples, while SOX10 and COL15A1 were closer to an average expression of 50 pseudobulk counts in both instances (range: 16-112). We estimate, based on prior knowledge about neuronal DRG expression, that pseudobulk counts of 50 or less are close to or within the noise range for these particular datasets.

### Literature review with semi-systematic search

The following search string was used in Pubmed on 28 September 2023: ((inflammatory soup) OR (inflammation)) AND ((DRG) OR (nociceptors) OR (sensory neurons)) AND ((calcium imaging) OR (electrophys*)). It was designed to help retrieve as many past papers studying inflammation-induced peripheral sensitization in dissociated peripheral neuron cultures as possible. Abstract screening and data extraction was performed by our three first authors (AL, YL & LF). We included only studies that examined DRG or trigeminal ganglion neurons in response to an inflammatory agent or solution with either patch clamping or calcium imaging. Moreover, the work, including the neuronal sensitization step, had to have taken place *in vitro*, e.g. we excluded experiments in which an inflammogen was injected into a mouse and differential excitability was subsequently investigated in dissociated cultures. The following information was extracted from papers which were included: which species was used and whether the cultures were dissociated DRG or TG; what pro-inflammatory stimulus was used; how long the treatment was applied; how many cells/coverslips were studied; what changes were reported in terms of patch clamp and calcium imaging parameters; and whether we could observe clear shortcomings in terms of design, reporting and/or sample size. Raw results are provided in **Supplementary Table 3**.

## Results

### Characterisation of iPSC-derived sensory neurons (iSNs)

We derived sensory neurons from two healthy human iPSC lines: Kute4 and UKB. UKB was modified from its originally founder line UKBi013-A to stably express the calcium sensor GCamP6f. Following a well-established protocol [7], both lines were differentiated into sensory neuron cultures with high expression of sensory neuron marker BRNA3A and neuronal marker PGP9.5 **(Figure 1)**. There was batch-to-batch variation in the percentage of BRNA3A+ cells, but wells of highly pure (90%+) neurons could be generated in the majority of differentiations. Depending on the requirements of particular downstream experiments, more or less pure wells were chosen based on assessment of cultures via light microscopy.

**Figure 1:**
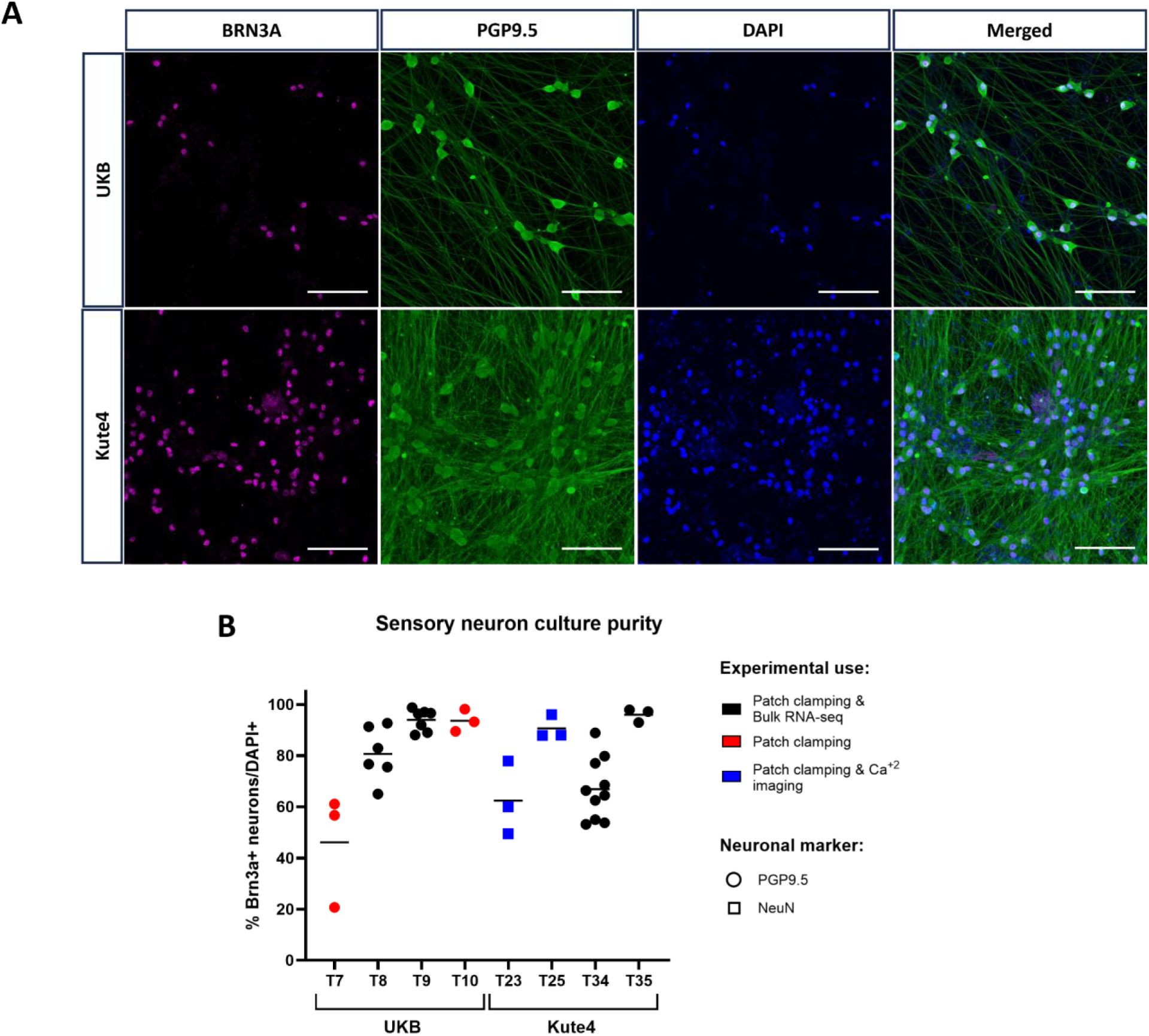
Immunostaining analysis demonstrates that pure sensory neurons could be differentiated from both our stem cell lines. (A) Representative images of UKB (day 54) and Kute4 (day 48) iPSC-derived sensory neurons. Scale bars: 100µm (B) Percentage of pure iSNs was determined for neurons from each independent differentiation used in subsequent experiments. Neurons aged day 46-81 were fixed and stained. Purity of iSNs was measured by taking the ratio of cells double positive for BRN3A and PGP9.5 or NeuN over DAPI+ nuclei. Each dot represents an individual coverslip. IDs T7-T35 refer to individual differentiations starting with UKB or Kute4 iPSC lines respectively.

As intended, calcium signals could be picked up directly in UKB-derived sensory neurons, with 50mM KCl-induced depolarisation clearly visible when cells were excited at 488 nm **(Figure 2).** Moreover, when neurons were stimulated with the sodium channel modulator veratridine, the fast dynamics of GCaMP6f enabled us to observe the expected and distinct response patterns that have previously been described in mouse DRG neurons [27] (**Figure 3** & **Supplementary Video 1**). We thus anticipate that the UKB iPSC line will be a useful tool for the functional study of human sensory neurons.

**Figure 2:**
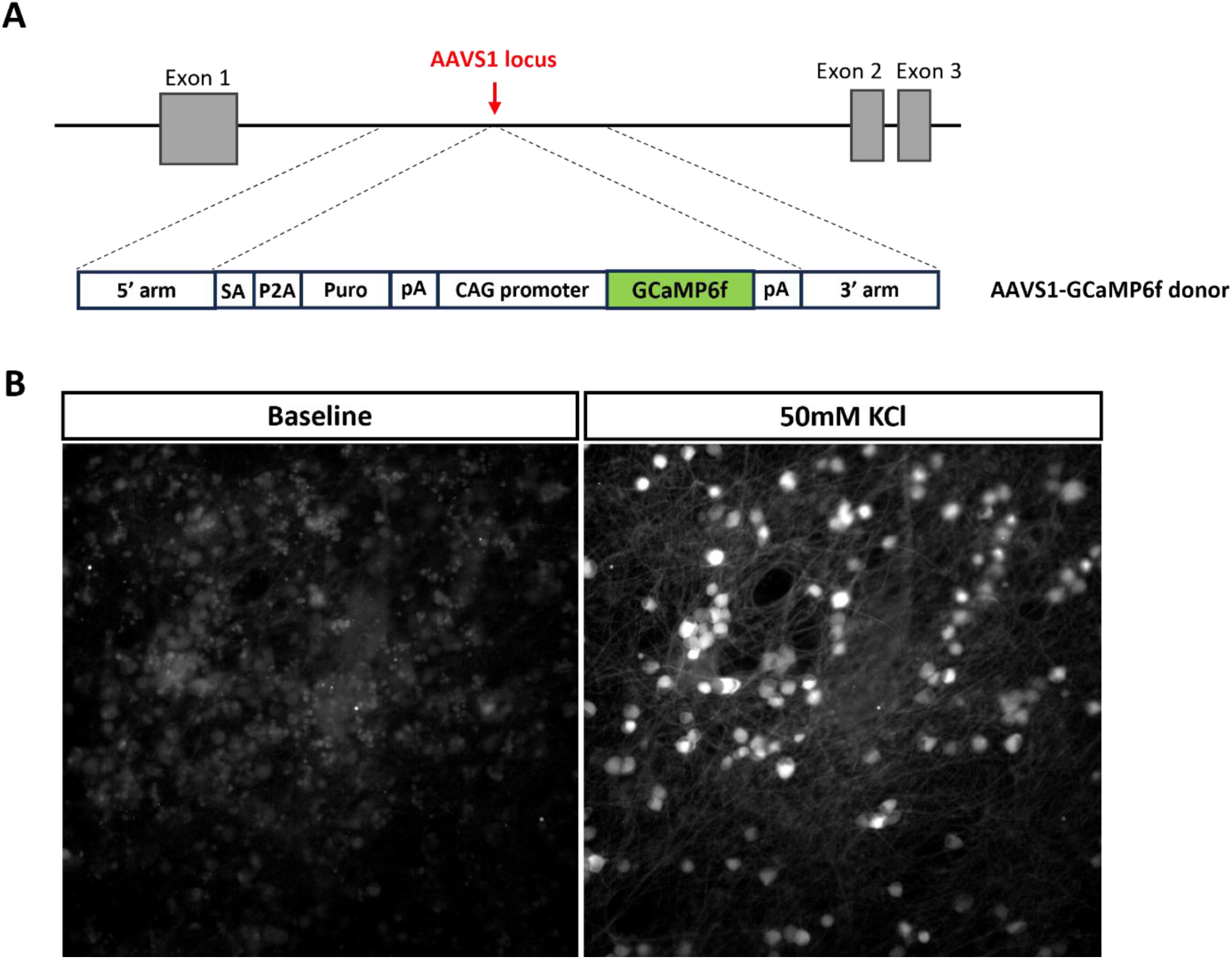
A novel iPSC line was generated which constitutively expresses GCaMP6f. (A) Schematic overview of the genomic insertion of GCaMP6f in the AAVS1 safe harbour locus of the UKB line. The insert contained GCamP6f driven by the CAG promoter, as well as 3’ and 5’ homology arms, a 3’ splice acceptor site (SA), a P2A self-cleaving peptide sequence (P2A) and a puromycin resistance gene (puro) terminated by a poly-adenylation signal (pA). (B) Screenshots of UKB-derived iSNs during calcium imaging, illustrating GCaMP6f fluorescent signal before and after application of 50 mM KCl.

**Figure 3:**
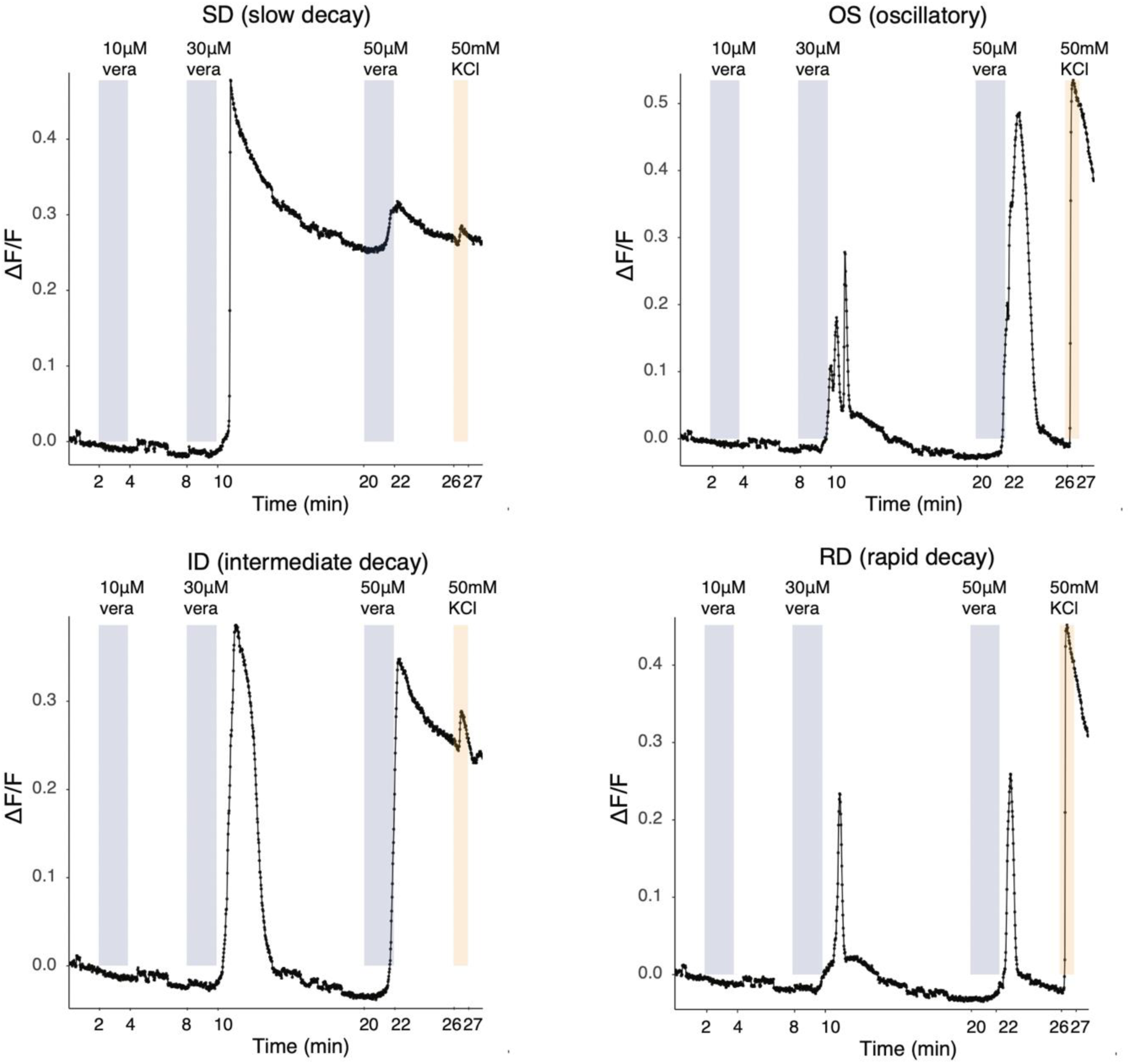
Representative traces of UKB-derived iSNs responding to veratridine demonstrating that the line’s endogenous GCamP6f can be used to detect fast, dynamic responses. UKB iSNs respond to veratridine with distinct patterns, similar to mouse DRG neurons, as characterised by [27]. Neurons (82 days old) were subject to three different concentrations of veratridine. 50 mM KCl was used as a positive control.

To confirm the transcriptional identity of our iSNs and to better understand their maturation state, we bulk sequenced RNA from these neurons at day 29-30 (D30), day 50-53 (D50) and day 69-70 (D70) across four differentiations and two iPSC lines **(Figure 4 & Supplementary Table 2**). We found that iSNs across the three timepoints expressed common neuronal (TUBB3, PRPH), sensory neuron-specific (ISL1, POU4F1), peptidergic (TAC1) and non-peptidergic (SST, P2RX3) markers. Pluripotency marker genes (NANOG, POU5F1) were fully downregulated. Finally, gene expression of markers for non-neuronal cell types (COL15A1 for fibroblasts, FABP7 for satellite glia, SOX10 & MBP for Schwann cells) was low or absent, confirming good purity of iSNs.

**Figure 4:**
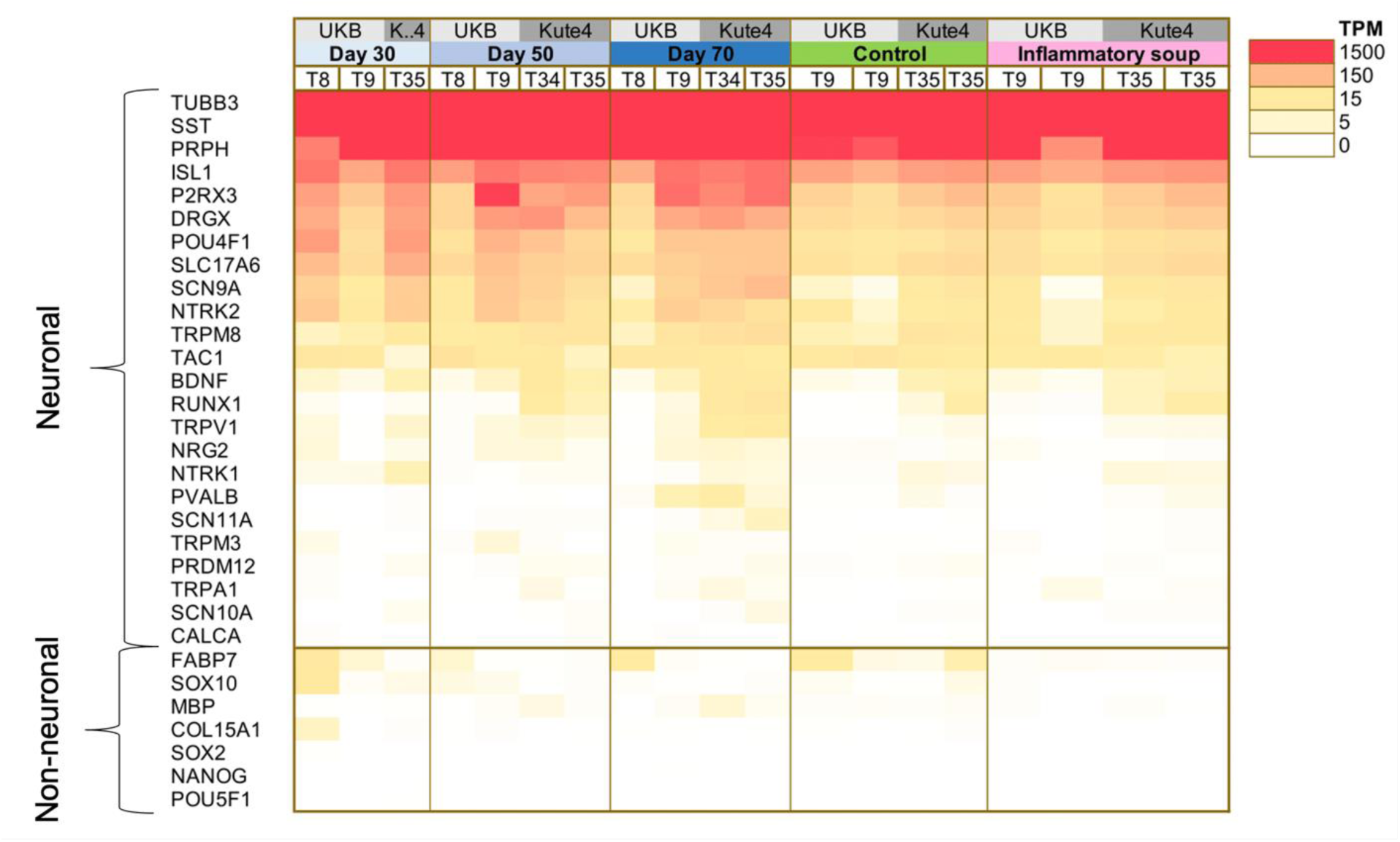
iSNs express common sensory neuronal markers, with low expression of contamination markers. Heatmap showing TPM expression values for selected neuronal and contamination markers. T8, T9, T34 and T35 are independent differentiations of two iPSC lines: UKB and Kute4, respectively. For the control and modified inflammatory soup conditions, cells were treated for 24h before RNA was extracted.

We were also able to confirm that our iSN model expresses receptors for several of the mediators present in our modified IS (**Figure 5**). NGFR (low-affinity NGF receptor), HTR1B & HTR1E (two of many serotonin receptors), TNFRSF1A (TNF receptor) and HRH3 (one of the histamine receptors) were clearly expressed above threshold in all samples. The expression of PTGER4 was at Transcripts per Million (TPM) > 1 in all but one sample, with an average of TPM of 3.9. NTRK1, the high affinity NGF receptor was expressed at TPM > 1 in 8 out of 16 samples. When we compared these data to pseudobulk expression levels derived from two independent single-nucleus RNA-seq datasets of human post-mortem DRG neurons [18; 29], we identified important commonalities and differences. NGFR, PTGER4, TNFRSF1A and HRH3 appeared comfortably expressed, similar to our iSN model system. However, in contrast to iSNs, HTR1E appeared more sporadically expressed and HTR1B seemed to be undetectable in all but 2 of 22 DRG neuron samples. Finally, NTRK1, PTGER3, HTR3A and HRH1 were present in post-mortem DRG samples, but not iSNs. Expression of these genes is highly likely to derive from neuronal nuclei, although it is impossible to categorically rule out contamination in the case of PTGER3, HTR3A and HRH1, due to the presence of glial and fibroblast-like transcripts in some of the post-mortem DRG samples. Publicly available sequencing repositories suggest that none of these transcripts are likely to be found in Schwann cells or satellite glia (e.g. [23] https://rna-seq-browser.herokuapp.com/), but PTGER3 and HRH1, for example, can be found in perineurial fibroblast-like cells, at least in mouse ([12] https://snat.ethz.ch/search.html?q=trpa1).

**Figure 5:**
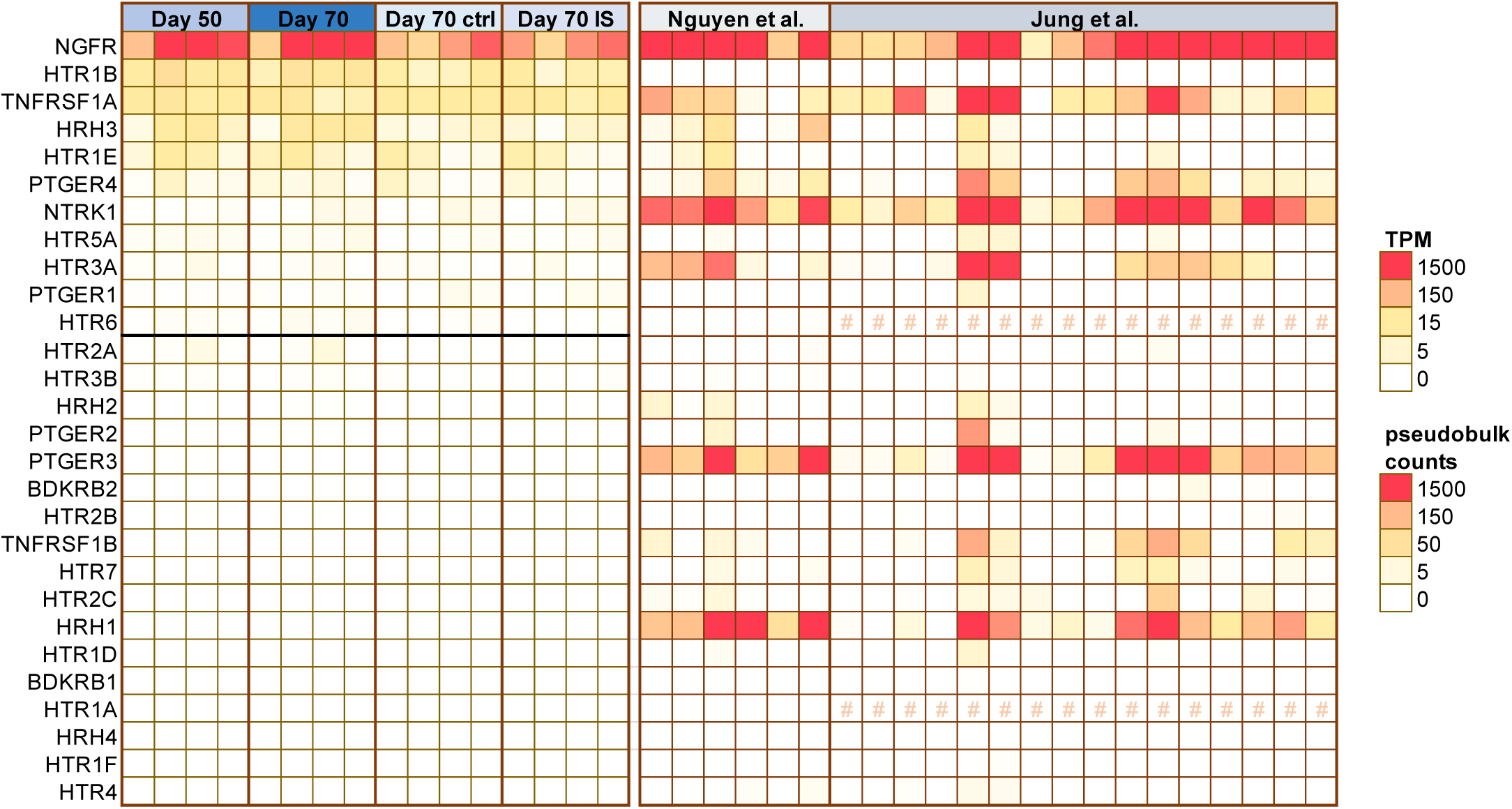
iSNs express receptors for several ligands within the modified IS. TPM expression values for our iSN RNA-seq data are shown on the left. Each square is a sample, arranged in the same order as in Figure 4. All genes above the black horizontal line are expressed at TPM > 1 in at least 7 samples or more. Genes below the line fall within the noise range of our particular dataset. On the right, we plotted pseudobulk counts derived from human postmortem DRG, specifically single-nucleus RNA-seq data published by two independent groups (Nguyen et al. [29] and Jung et al. [18]). Each square displays data on DRG neurons obtained from a different individual (n = 22 postmortem samples). We estimate that counts less than 50 are within the noise range. HTR1A and HTR6 were not annotated within Jung et al. and are therefore labelled with an #.

When comparing across timepoints, we found that D30 iSNs were the most distinct, with D50 and D70 samples clustering together (**Figure 6**). Indeed, when comparing pairs of samples, D70 vs. D30 had the highest number (949) of differentially expressed genes, followed by D50 vs. D30 (312). Noticeably, only 23 genes were differentially expressed between D70 vs. D50 (**Supplementary Table 2**). They include genes like PVALB and CDH8 (more abundant at D70) and BACH1 (more abundant at D50), suggesting some difference in sensory neuron phenotypes between the two groups. It is unclear whether this is a systematic, replicable difference denoting a further increase in maturity at D70, or whether it simply reflects the variability expected from small molecule iSN differentiations. Meanwhile, neurons at D30 are obviously different, with clear signs of immaturity, e.g. lower expression of homeobox D1 (HOXD1), a known NGF-sensitive regulator of nociceptor circuitry in invertebrates [13], and limb-expressing 1 (LIX1), which has been linked to sensory neuron and nociceptor development. Specifically, Lix1 is reported to be significantly downregulated in Brna3-null [10] and Islet-1 knock-out mice [42], both of which lack sensory neuron innervation.

**Figure 6:**
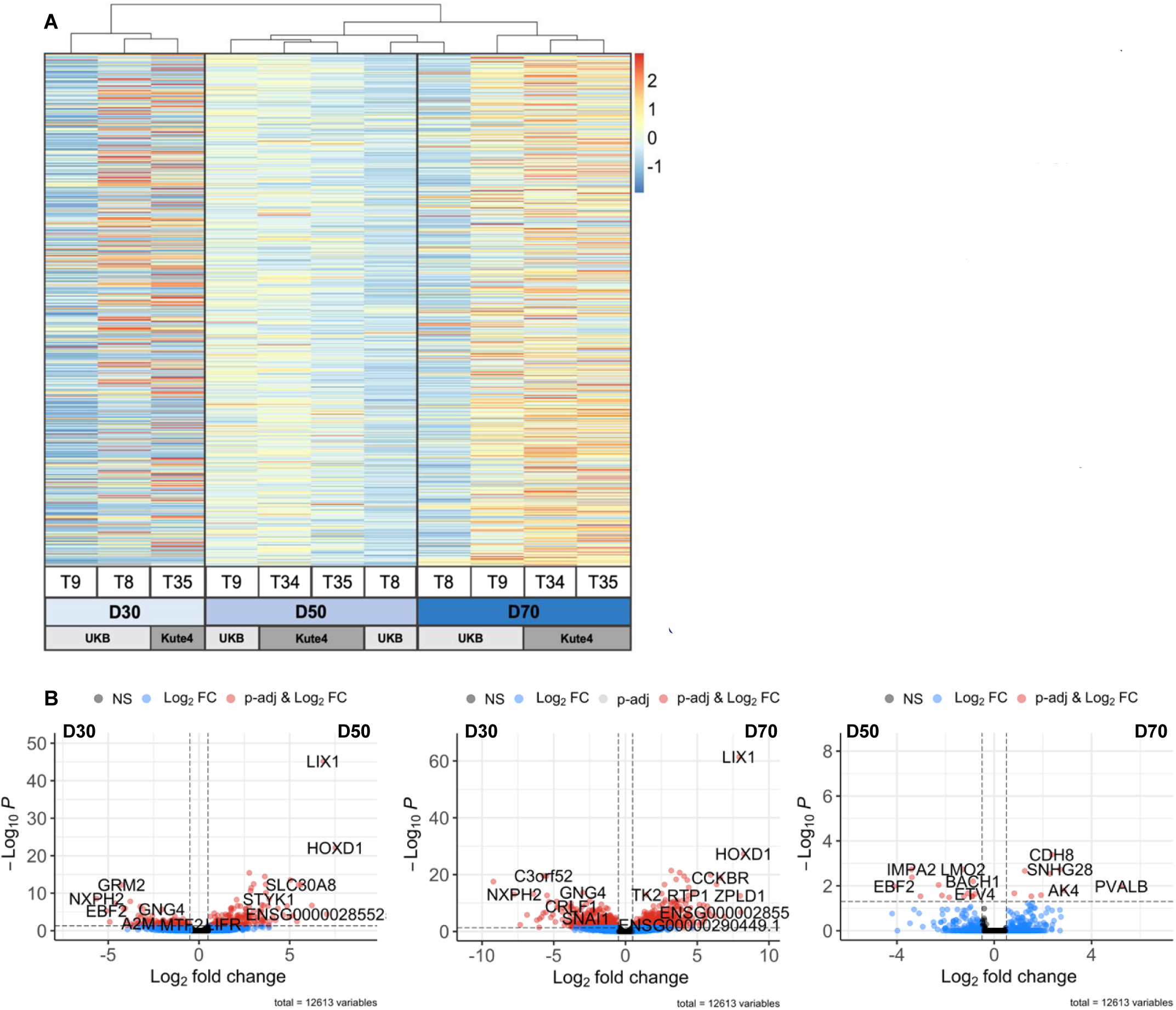
iSNs reach a consistent level of maturity beyond differentiation day 50. (A) Heatmap displaying differentially regulated genes across three iSN differentiation timepoints (D30, D50 and D70). Colours represent z-scored TPM values (by gene). Columns represent individual samples and were ordered using unsupervised clustering with the hclust function (method = ward.D2). Clustering was performed on all genes which were expressed according to our cut-off (see Methods), differentially regulated at adj. p < 0.05 and regulated at log2FC > 1.5 or log2FC < -1.5. (B) Volcano plots displaying differentially expressed genes for each time point comparison. Adjusted p cut-off: <0.05, Log2FC cut-off: > 0.5 or < -0.5. Plots created using Enhanced Volcano R package [2]. T = trial indicating independent differentiations of Kute4 and UKB lines.

### Functional analysis of iSNs in response to modified inflammatory soup

We next assessed the effects of modified-IS on sensitising iSNs via calcium imaging. Following incubation in the soup for 24h, neurons (day 54-74) were subjected to 50 μM veratridine, which opens voltage-gated sodium channels or 100 μM pregnenolone sulphate, which activates TRPM3 channels, a receptor that may convey information about noxious heat [44]. In neither case could we observe a difference in the percentage of responding cells or in the magnitude of responses (**Figure 7** & **Supplementary Figure 1**). Neurons showed similar response dynamics over the course of the experiments. Notably, this result did not seem to be affected by the presence of non-neuronal cells: we noted that one trial (T23) was less pure than the other (T25) (see **Figure 1B**), however, both showed similar responses (grey vs. orange dots in **Figure 7D&E).**

**Figure 7.**
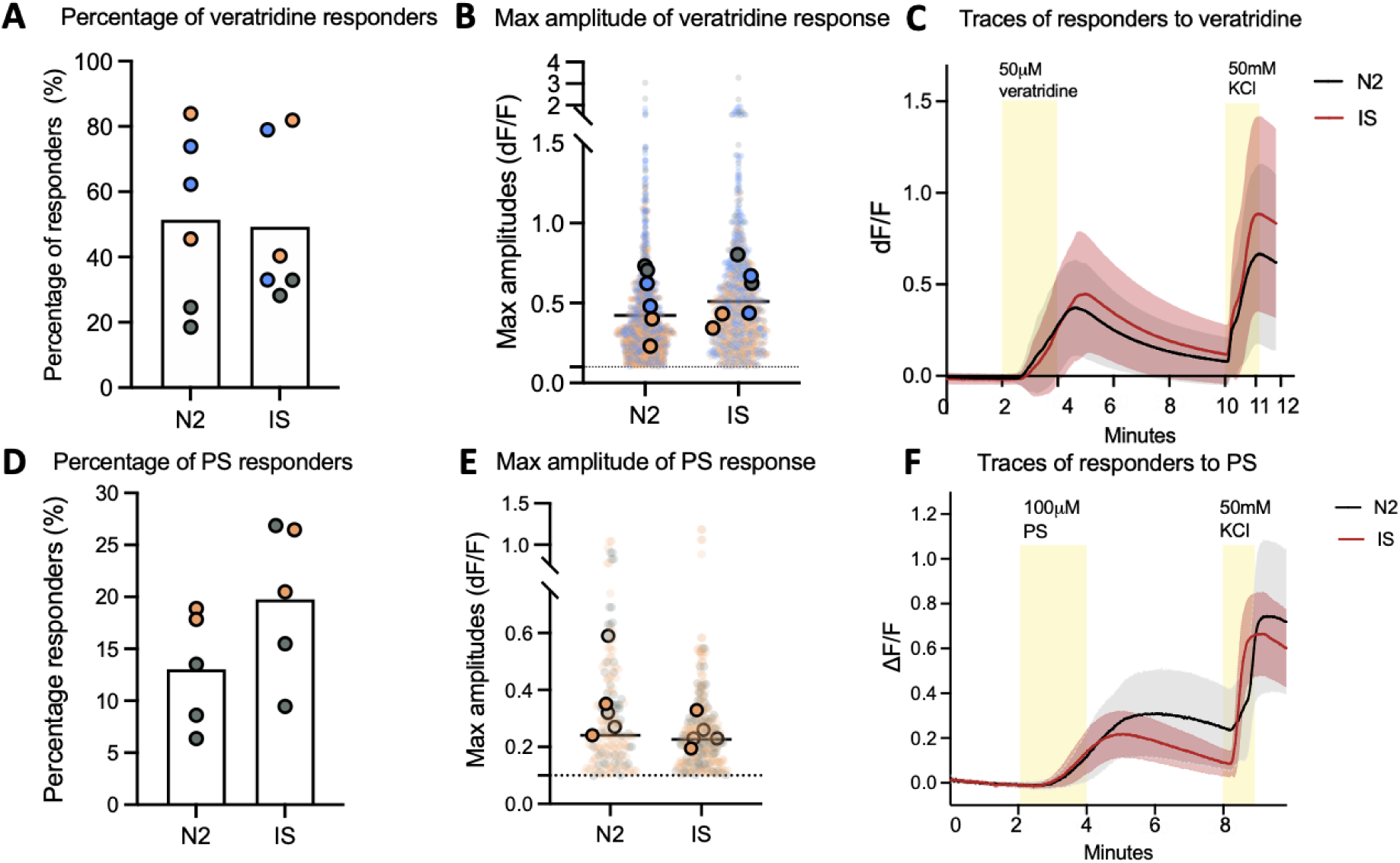
Modified-IS does not sensitise Kute4-derived iSNs in response to veratridine or pregnenolone sulphate (PS). The inflammatory soup does not change the response percentage (**A,D**) or the maximal amplitude of response (**B,E**) to 50 μM veratridine or 100 μM pregnenolone sulphate (PS). Each big dot in **A & D** represents a coverslip. In **B& E**, the small, transparent dots are the maximal values of each cell in each coverslip. The superimposed large-sized dots are the mean values of each experiment. The dotted line is at 0.1, the cut-off for responders. **C & F** plot average traces in response to modified-IS and N2 control medium over the course of the experiment; the shades surrounding the average line are SD at each time point. For each condition, the trace is the average of 3 experiments. See **Supplementary Figure 1** for individual representative traces. For all experiments, neurons were incubated in N2 (media control) or modified-IS for 24 hours before the experiment. Recordings took place at room temperature. Neurons were differentiated from Kute4 iPSC for at least 50 days. Different colours indicate independent differentiations (T23-29); for veratridine: grey = T28 72 days old; orange: T29 57 days old; blue: T29 62 days old; for PS experiments: grey = T25 63 days old; orange = T23 74 days old.

Traditionally, peripheral sensitization in a dish was most often assessed using direct measures of neuronal activity, rather than with indirect tools like calcium imaging. We therefore undertook whole-cell current-clamp recordings of iSNs in control conditions (Ctrl) or treated with modified-IS for 24 hours (**Figure 8)**. We had previously been involved in an effort where such recordings revealed increased excitability in mouse DRG nociceptors treated with conditioned medium from activated fibroblasts [6]. This medium was shown to contain a cocktail of cytokines and pro-inflammatory molecules, though none that are within the modified-IS we used here. Nevertheless, we found that treating iSNs with modified-IS resulted in more depolarised resting membrane potential compared to the control group (two tailed unpaired t-test, *p=0.0383*, Ctrl: -71.31 ± 1.1 mV; IS: -67.74 ±1.29 mV). In contrast, modified-IS did not have any effect on passive membrane properties of iSNs, with capacitance and input resistance very similar between the two groups. To study whether inflammatory soup changed any neuronal firing properties we also assessed spontaneous action potential firing, rheobase, action potential firing characteristics and frequency upon increasing current injection. The neurons from both groups behaved in a similar way regardless of the treatment. Finally, modified-IS did not result in any changes in evoked action potential properties, evident by the analyses of: peak amplitude, time to peak, action potential threshold, half-width, fast afterhyperpolarisation (fAHP) and medium afterhyperpolarisation (mAHP).

**Figure 8.**
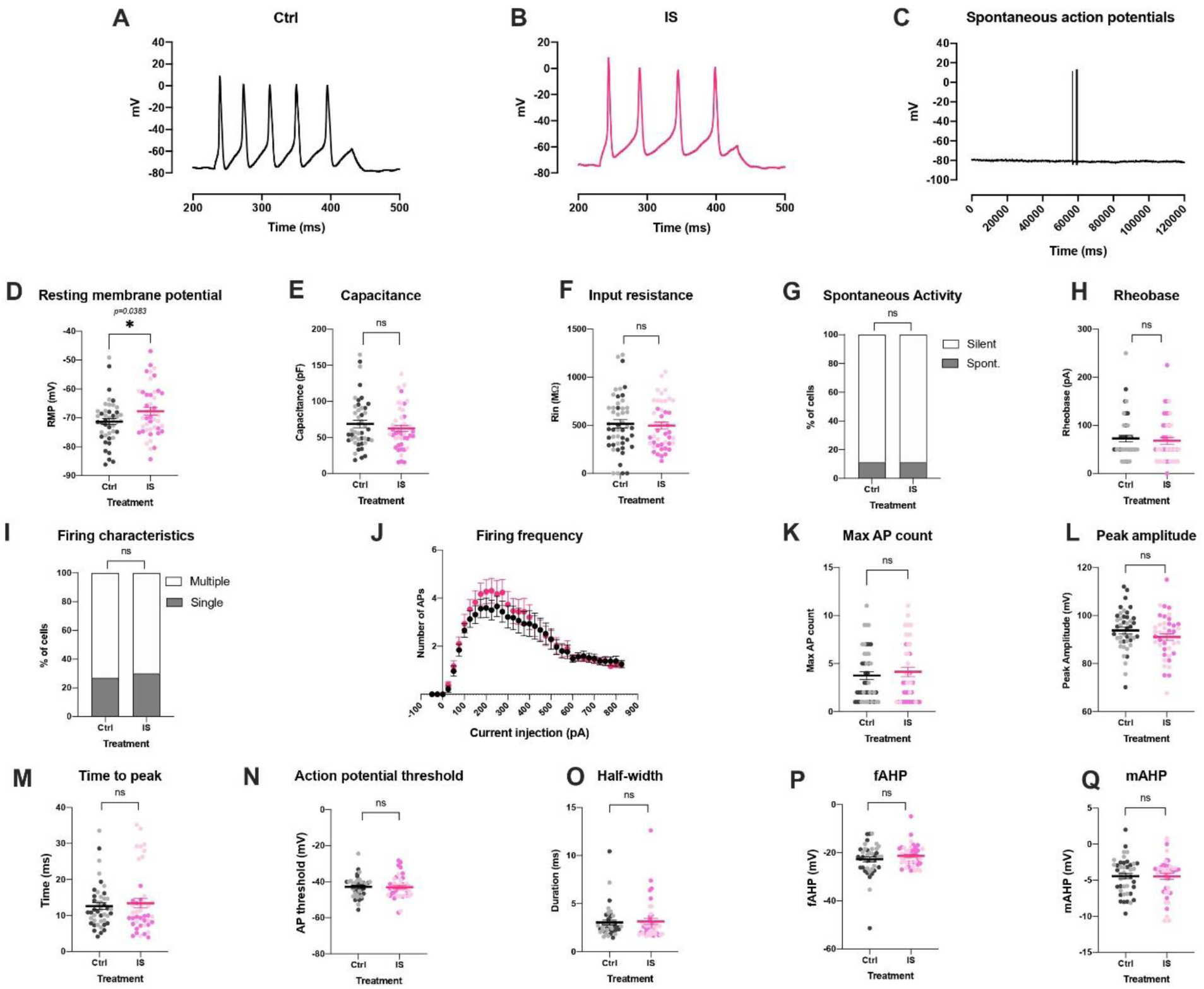
Modified-IS induces only a very minor change in resting membrane potential in iSNs, while all other electrophysiological properties of the neurons appear unaffected. Representative traces of evoked action potentials in control (A) and inflammatory (B) conditions and of spontaneous action potentials (C). (D) The resting membrane was more depolarised in modified-IS treated compared to control neurons (two tailed unpaired t-test, p=0.0383). Modified-IS did not elicit any differences on: capacitance (E), input resistance (F), spontaneous activity (G), rheobase (H), firing characteristics (I), firing frequency upon current injection (J), number of maximal action potentials (K), peak amplitude (L), time to peak (M), action potential threshold (N), half-width (O), fast afterhyperpolarisation (fAHP) (P) and medium afterhyperpolarisation (mAHP) (Q). Data were tested for normality, two-tailed unpaired t-tests or Mann– Whitney tests were used accordingly for: D, E, F, H, L, M, N,O. Fisher’s exact test was used to assess any potential changes in spontaneous activity and firing characteristics. Repeated measures mixed effects analyses were used to assess any changes in firing frequency between the two groups (J) with Bonferroni posthoc test: current injection resulted in significant changes in firing frequency (p<0.0001), however, neither treatment (Ctrl vs IS) nor current injection x treatment was significant. All data represent mean ± SEM pooled from five different differentiations per line, n=45 per treatment group. Darker colour represents Kute4 line, lighter colour UKB line. Ctrl: Kute4 n=22, UKB n=23; modified-IS: Kute4 n=21, UKB n=24. * p<0.05

These patch clamp data were generated using our two different iPSC lines over five independent differentiations. All neurons were at least 45 days old but varied in age range from 52-63 Kute4 and Day 45-55 UKB. Yet, we could not detect any discernible patterns or differences between differentiations **(Supplementary Figure 2)**. We also examined whether the presence of non-neuronal cells might have impacted on neuronal excitability. For this, we grouped our iSN batches based on their purity as determined in **Figure 1** and analysed our functional data accordingly. Differentiation batches were defined to be impure if the mean % of pure iSNs was below 70%, as such, T7, T23 and T34 were defined as “*impure”* and T8, T9, T10, T25 and T35 as “*pure*”. We specifically focused on electrophysiological parameters that have been identified by us or others in the literature to be modified when DRG neurons are treated with inflammatory stimuli (**Figure 9**). A few differences could be observed at face value, specifically a more prominent increase in resting membrane potential in pure trials and a more obvious difference in spontaneous activity in impure trials. However, it is unclear whether these would be replicable, given that our analyses were restricted to a low number of neurons in the impure group (n = 9 in ctrl and n = 6 in modified-IS).

**Figure 9.**
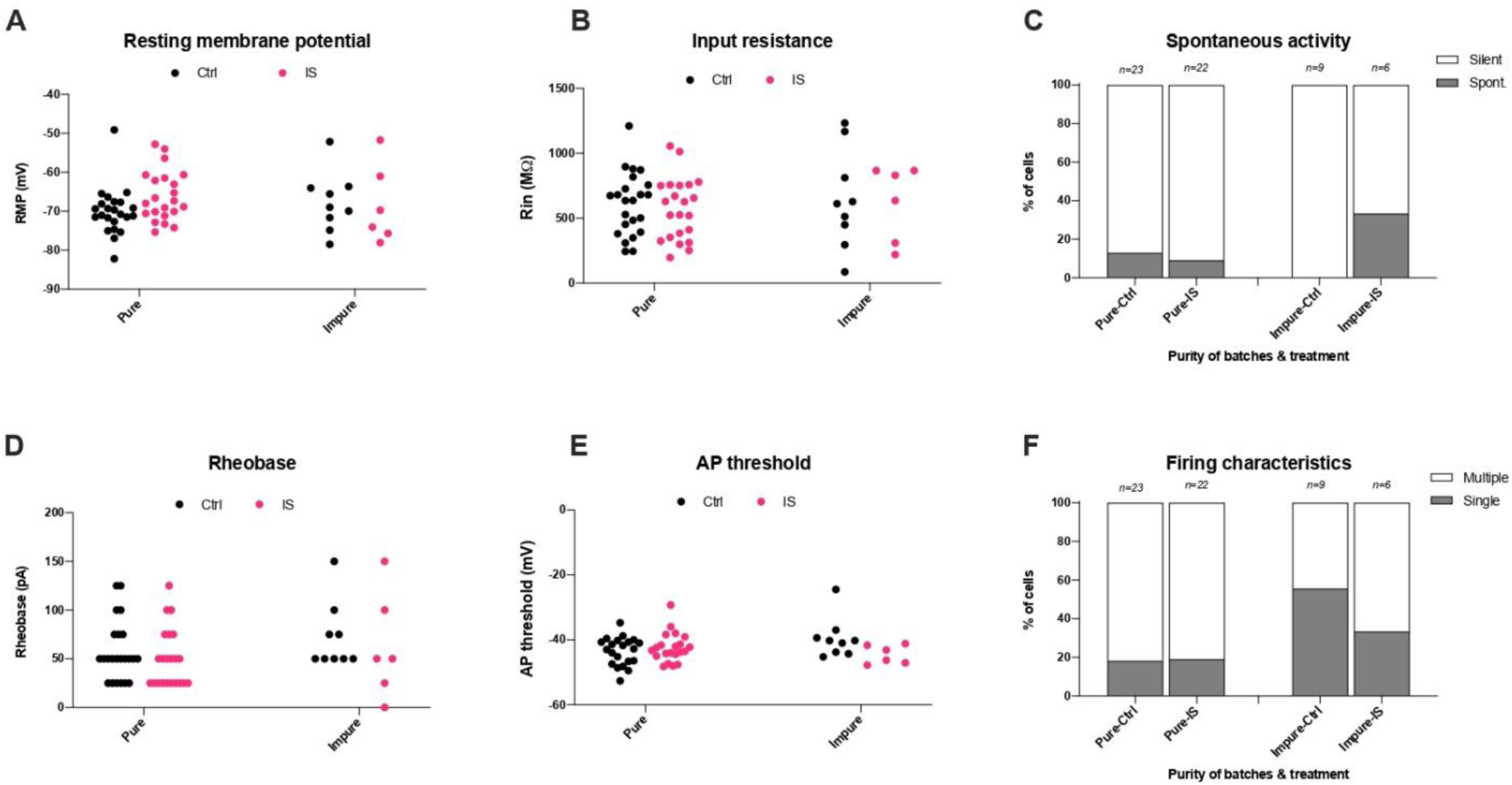
Differentiation batches which are less pure are not more likely to show signs of peripheral sensitisation. Comparison of the effect of modified inflammatory soup on electrophysiological parameters reported from the literature between “pure” and “impure” iSNs. (A) Resting membrane potential, (B) Input resistance, (C) Spontaneous activity, (D) Rheobase, (E) Action potential threshold, (F) Firing characteristics. Pure trials (T8, T9, T10, T25 and T35), Ctrl n=23, mod. IS n=22; Impure trials (T7, T23 and T34), Ctrl n=9, mod. IS n=6.

### Transcriptional changes in response to modified inflammatory soup

Following our functional experiments using 24-hour incubation with modified-IS, we sought to investigate whether iSNs change transcriptionally under these conditions (**Figure 10**). We used D60 iSNs from two differentiations, one from each iPSC line. Perhaps unsurprisingly, given the relative functional similarity between neurons with and without inflammatory mediators, a principal component analysis (PCA) revealed samples to cluster by iPSC identity rather than treatment condition. Furthermore, differential expression analysis showed only one gene (MIA) significantly regulated at adj. p < 0.05, illustrating near zero transcriptional changes in response to modified IS. No additional changes were revealed when each iPSC line was analysed individually (**Supplementary Figure 3**).

**Figure 10:**
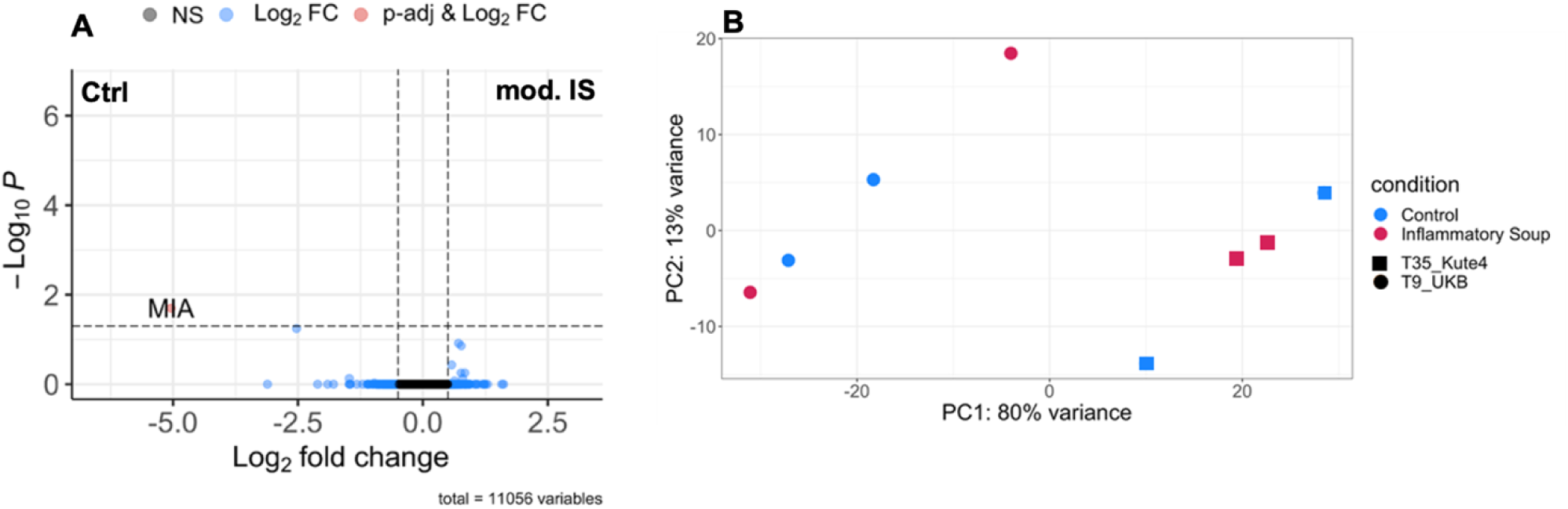
Incubation with modified IS does not alter the transcriptome of iSNs. (A) Volcano plot displaying differentially expressed genes. Adjusted p cut-off: <0.05, Log2FC cut-off: > 0.5 or < -0.5. (B) PCA plots displaying points coloured by condition (control in blue and modified IS in red) and shaped to indicate different iPSC lines (square for Kute4, circle for UKB).

### Functional analysis of mouse DRG cultures in response to modified inflammatory soup

Given that we observed only very modest effects in our iSN when treated with modified-IS, we wanted to confirm whether this could have been due to limitations of our stem-cell derived model system. It is well-established that sensory neurons differentiated with the Chambers protocol, like ours, lack receptors known to be important for peripheral sensitization, such as TRPV1 and the tetrodotoxin-resistant sodium channel SCN10A (**Figure 4**). Mouse neurons treated with modified-IS for 24 hours displayed a small, but non-significant increase in resting membrane potential (**Figure 11**). There was no effect on cell capacitance, but a reduction of input resistance (Ctrl: 839.35 ± 88.4 MW; IS: 561.967 ± 64.94 MW, two-tailed Mann–Whitney test). One might speculate that reduction of input resistance without affecting cell capacitance denotes an increased number of open channels, possibly as a result of the action of inflammatory mediators on neurons.

**Figure 11.**
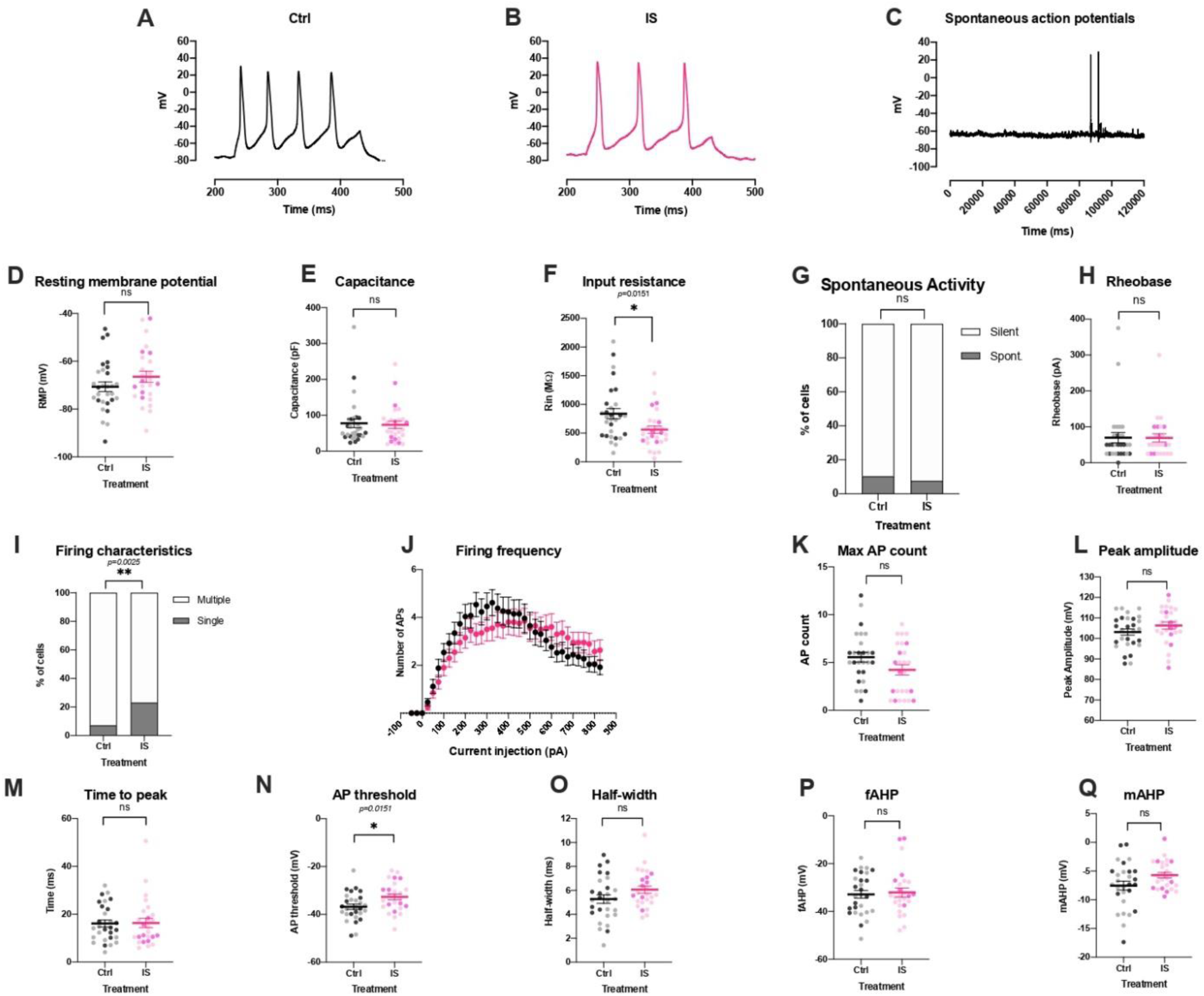
Modified-IS induces only very few, modest changes in the electrophysiological properties of mouse DRG neurons. Representative traces of evoked action potentials in control (A) and inflammatory (B) conditions and of spontaneous action potentials (C). mod-IS resulted in significant changes in input resistance (F) (Mann–Whitney test, p= 0.0153), firing characteristics (I) (Fisher’s exact test, p=0.0025) and action potential threshold (N) (unpaired t-test p=0.0151). IS did not elicit any differences on: resting membrane potential (D), capacitance (E), spontaneous activity (G), rheobase (H), firing frequency upon current injection (J), number of maximal action potentials (K), peak amplitude (L), time to peak (M), half-width (O), fast afterhyperpolarisation (fAHP) (P) and medium afterhyperpolarisation (mAHP) (Q). Data were tested for normality, and unpaired t-test or Mann–Whitney tests were used for (D, E, F, H, L, M, N, O). Fisher’s exact test was used to assess any potential changes in spontaneous activity and firing characteristics. Repeated measures mixed effects analyses were used to assess any changes in firing frequency between the two groups (J) with Bonferroni posthoc test: current injection resulted in significant changes in firing frequency (p<0.0001) and current injection x Treatment (p=0.0002), however, treatment alone (Ctrl vs IS) did not elicit any significant differences. All data represent mean ± SEM. DRG neurons were patched following 5-7 days in culture (Ctrl n=26; IS n=20) and following 3 days in culture (Ctrl: n=4, IS n=6), total Ctrl: n=29, IS: n=26. Darker colour represents cells isolated from male mice, lighter colour from female mice. * p<0.05; ** p<0.01.

In terms of neuronal firing properties, modified-IS did not result in significant changes in the percentage of mouse neurons firing spontaneous action potentials nor changes in rheobase or firing frequency upon current injection. However, we found that an increased number of cells in the modified-IS group fired single action potentials compared to control (two-sided, Fisher exact test *p=0.0025, Ctrl n=29; IS n=26*). We analysed the action potential shape and found that modified-IS resulted in a significantly more depolarised action potential threshold (Ctrl: -32.7 ± 2.58 mV; IS: -36.83 ± 1.11 mV, two-tailed unpaired t-test, *p=0.0151*). However, none of the other parameters related to action potential shape were significantly different, including peak amplitude, time to peak, half-width, fast afterhyperpolarisation (fAHP) and medium afterhyperpolarisation (mAHP). Moreover, more single action potentials and an increased action potential threshold would indicate decreased excitability rather than the increased excitability one would predict with peripheral sensitization.

### Review of the literature using a systematic search and qualitative analysis

Finally, to aid interpretation of our data, we decided to compare our results to what has been reported previously in the literature. We undertook a semi-systematic search in Pubmed and screened 336 articles of which 59 were included in a data extraction step. Upon full-text screening a further 8 were excluded because they did not use an inflammatory substance to treat the cells, while a further 12 were excluded because they only recorded specific channel currents rather than reporting on the more general action potential parameters we measured here. This left 39 studies from which we obtained qualitative data. The most commonly used pro-inflammatory treatments were PGE2 (n = 5) or other COX2-pathway related substances (n = 3), TNF (n = 4) and pro-inflammatory supernatants or conditioned media (n = 6). Three studies examined inflammatory soup either in its original form (n = 2) or in a modification designed to mimic activated mast cells (n = 1).

In line with our results, there seemed to be little consensus on exactly how pro-inflammatory factors affect sensory neuron excitability *in vitro*. Very few studies examined spontaneous activity (2 out of the 21 which conducted patch clamping). By far the most common alterations to be reported (18/21, i.e. 86%) were related to neuronal firing properties, with 2 studies reporting the same increase in action potential threshold that we observed. But there were also just as many studies that reported a decrease in action potential threshold, and yet more that saw changes in peak amplitude, the number of spikes, or the latency to the first action potential. An increase in resting membrane potential was the most consistently observed change, reported in 7 out 21 papers (i.e. 33%). 22 papers published calcium imaging data, 11 of which reported changes in amplitude, 5 of which observed changes in the number of responders, 4 of which changes in both, and 2 no changes at all. Finally, the great majority of studies had significant uncertainty associated with them, with only 10 papers deemed without clear limitations around experimental design, reporting and/or sample sizes. For example, very low sample sizes were common, with only 4-9 neurons patched per condition. All relevant references and data associated with the search are provided in **Supplementary Table 3**.

## Discussion

This study set out to examine the response of human stem-cell derived sensory neurons (iSNs) to modified inflammatory soup (IS). We observed no significant changes in calcium dynamics or transcriptional responses, and only a very modest increase in resting membrane potential upon patch clamping.

There are several potential explanations for our results, which we will discuss in turn. Firstly, *in vivo*, it is likely that non-neuronal cells play an important role in bringing about peripheral neuron sensitization, not just as sources of IS-components, but also as intermediary cells which may release additional pro-inflammatory mediators that bind nociceptors. For example, given known mRNA receptor expression patterns, it is likely that bradykinin exerts at least some, if not all, of its pro-algesic effects via actions on e.g. keratinocytes [26]. To test for the role of non-neuronal cells, we compared the response of our human iSNs between ‘purer’ differentiations and those that were more contaminated with non-neuronal cells. We also performed patch clamping in conventional rodent DRG neuron cultures, dissociations of which contain a high proportion of non-neuronal cells [43], mostly made up of fibroblasts and a de-differentiated mix of Schwann and satellite glial cells [16]. We did not find any evidence that the presence of non-neuronal cells dramatically increases the extent to which sensory neurons respond to modified-IS in our dissociated *in vitro* settings.

As a second possible cause for the lack of sensitization, we focused on the inherent limitations of our iSN model system. Stem-cell derived sensory neurons are a great tool for the study of human peripheral nervous system function, but current differentiation protocols (the most commonly used of which is Chambers et al [7]) fail to generate the full range of neuronal sub-classes known to exist *in vivo*. In fact, rather than splitting into clear populations of A-fibres and peptidergic and non-peptidergic C-fibres, iSNs tend to form one single homogeneous group of neurons that express some markers that usually do not co-occur (e.g. SST and NF200) while lacking others, like TRPV1 and Na_v_1.8. The absence of the latter two may be particularly important in the context of peripheral sensitization, since phosphorylation of TrpV1 [28], followed by regulation of TTX-resistant sodium channels like Na_v_1.8 [1], is generally described as one of the key biological underpinnings of peripheral sensitization. Yet, if our failure to detect abnormal excitability in response to modified-IS was purely due to a quirk within our human model system, we would have expected to detect clear signs of peripheral sensitization when patching mouse sensory neurons. After all, the concept of peripheral sensitization is best studied in rodent models, with many of the original publications on inflammatory soup and its constituents using dissociated cultures [4; 20] or skin-nerve preparations for electrophysiological studies [19; 41].

Overall, in our electrophysiological experiments, we examined 14 different parameters, which is not at all unusual for patch clamp experiments of this kind. This is noteworthy: historically, multiple comparison corrections would not have been routinely applied to such analyses, even though, arguably, they are very necessary. Stringent correction methods, e.g. Bonferroni, would indicate that we should apply a p-value threshold of 0.003 to reduce the chances of false positive results. We observed only a single p-value that was below this threshold, with modified-IS inducing more single than multiple action potential spikes in mouse DRG neurons. Yet, this result is the opposite of what one would expect from neurons that are more excitable due to peripheral sensitization. All other ‘significant’ findings (increased resting membrane potential, reduced input resistance and increased action potential threshold) were sporadic and did not replicate across species. This is also a pattern that is repeated across the literature: our semi-systematic review indicates that past papers have reported a bewildering range of changes across varying patch clamp parameters, with no consistent differences emerging. This, coupled with the fact that single-cell electrophysiological techniques are inherently prone to unintentional p-hacking^1^, makes it more likely that some of the previously reported effects are false positives or, at least, inflated in size.

This then, is a third possible explanation for the results we observed, and currently the one we deem most probable: modified IS may simply not exert very large effects on the function of sensory neurons in primary mono-cultures – whether stem-cell derived or dissociated from animals. Given what we know about the substances within modified-IS, this is probably less of a reflection of the pro-algesic potential of individual components (shown to be high in *in vivo* settings) than a consequence of the highly artificial and reductionist environment of *in vitro* cell culture. The latter may lack key elements of an inflammatory environment, such as low pH or the presence of dysfunctional immune and/or stromal cells. Future studies may need to focus on increasing the effect size window in mono-cultures, e.g. by using assays that allow for long-term recordings and subsequent within-cell comparisons, like multi-electrode arrays. Further good options might be to use more complex, multi-cellular *in vitro* setups or other pro-inflammatory admixtures, e.g. those modelling the function (and mediator release) of specific pro-algesic cell subtypes, like activated fibroblast-like synoviocytes [6] or Marco+ macrophages [34].

In conclusion, here we have provided evidence to indicate that studying peripheral sensitization in dissociated cultures, whether human or murine in origin, is not straightforward. Even a mixture of very well-known pro-algesic mediators appears to induce only very modest functional effects in neurons, if any. More work is necessary to identify better-suited positive control substances and/or optimise cellular model designs to reliably demonstrate peripheral sensitization *in vitro*. We have also provided a couple of novel research tools that we hope will accelerate this and other efforts, specifically an iPSC line which constitutively expresses the calcium sensor GCamP6f and an R-based script to analyse its activity.

## Acknowledgements

AL was supported by the UK Medical Research Council (MR/N013700/1) and King’s College London as a member of the MRC Doctoral Training Partnership in Biomedical Sciences. YL and MS were funded by the Wellcome Trust as part of the Neuro-Immune Interactions in Health & Disease Wellcome Trust PhD Programme (218452/Z/19/Z). FD, LT and IZ were funded by a Wellcome Trust Collaborative Award (224257/Z/21/Z). LF was funded by an Ono Pharmaceuticals Rising Stars Award held by FD. AS was funded by FNRS and a Sofina Boël mobility grant.

The GCamP6f line was generated as part of an Innovative Medicines Initiative 2 Joint Undertaking under Grant Agreement no. 116072. This Joint Undertaking has received the support from the European Union’s Horizon 2020 research and innovation.

We acknowledge King’s College London as the source of HPSI0714i-kute_4 human iPSC line which was generated under the Human Induced Pluripotent Stem Cell Initiative funded by a grant from the Wellcome Trust and Medical Research Council, supported by the Wellcome Trust (WT098051) and the NIHR/Wellcome Trust Clinical Research Facility, and acknowledges Life Science Technologies Corporation as the provider of Cytotune.

This research was funded in whole or in part by UK Research & Innovation and the Wellcome Trust. For the purpose of Open Access, the author has applied a CC BY public copyright licence to any Author Accepted Manuscript (AAM) version arising from this submission.

OB is a cofounder, CEO and shareholder of LIFE & BRAIN GmbH. None of the other authors have any conflicts of interest to declare in relation to this work.

## Supplementary Figures

**Supplementary Figure 1.**
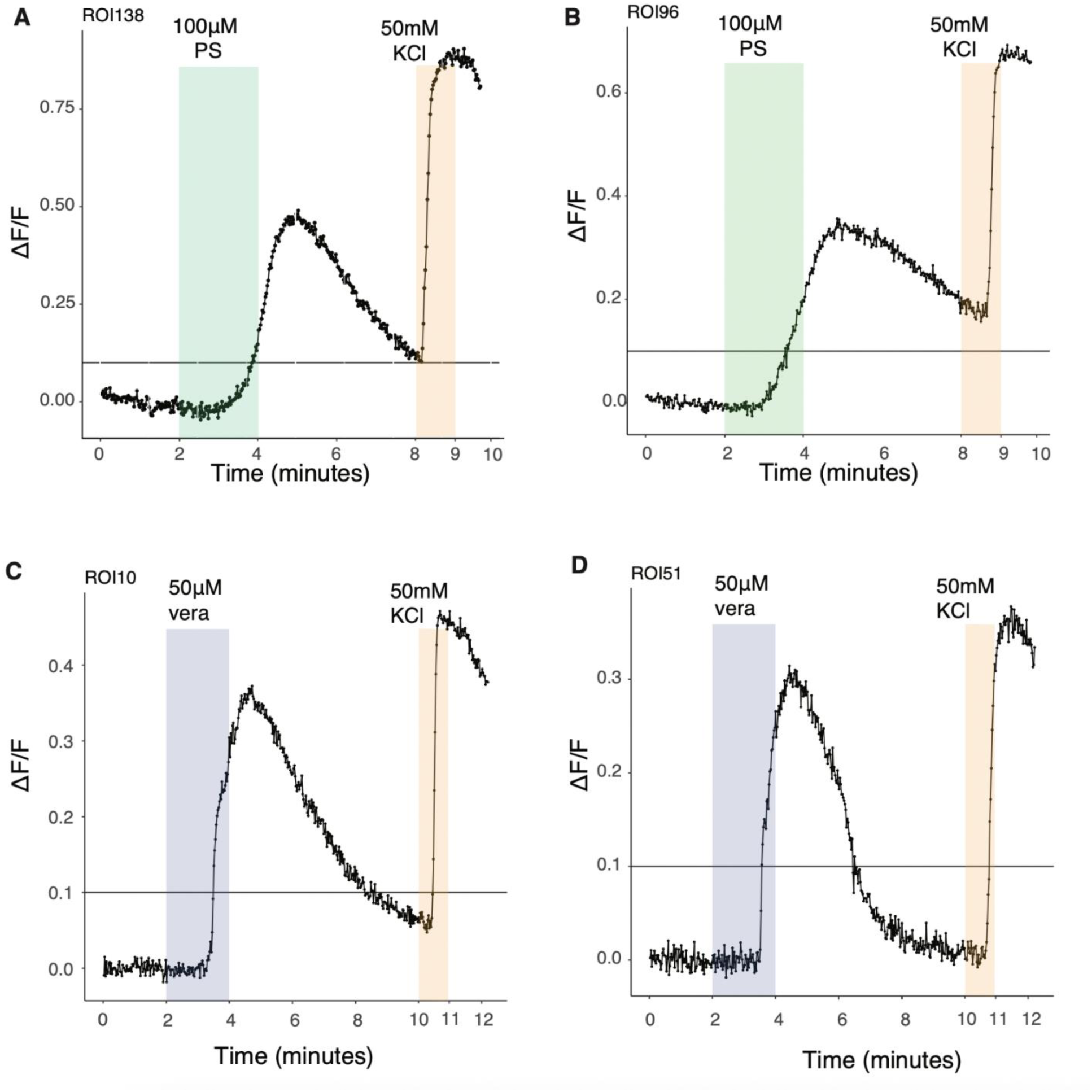
Example raw traces of neuronal response to 100μM PS and 50μM veratridine. 50mM KCl was used as a positive control at the end of the experiment. Example responder traces of 100μM PS for neurons incubated with IS (A) or incubated with N2 (B) for 24h (Kute4 iPSC line, trial T25, differentiated until day D63). Example responder traces of 50μM veratridine for neurons incubated with IS (C) or incubated with N2 (D) for 24h (Kute4, T29, D54).

**Supplementary Figure 2.**
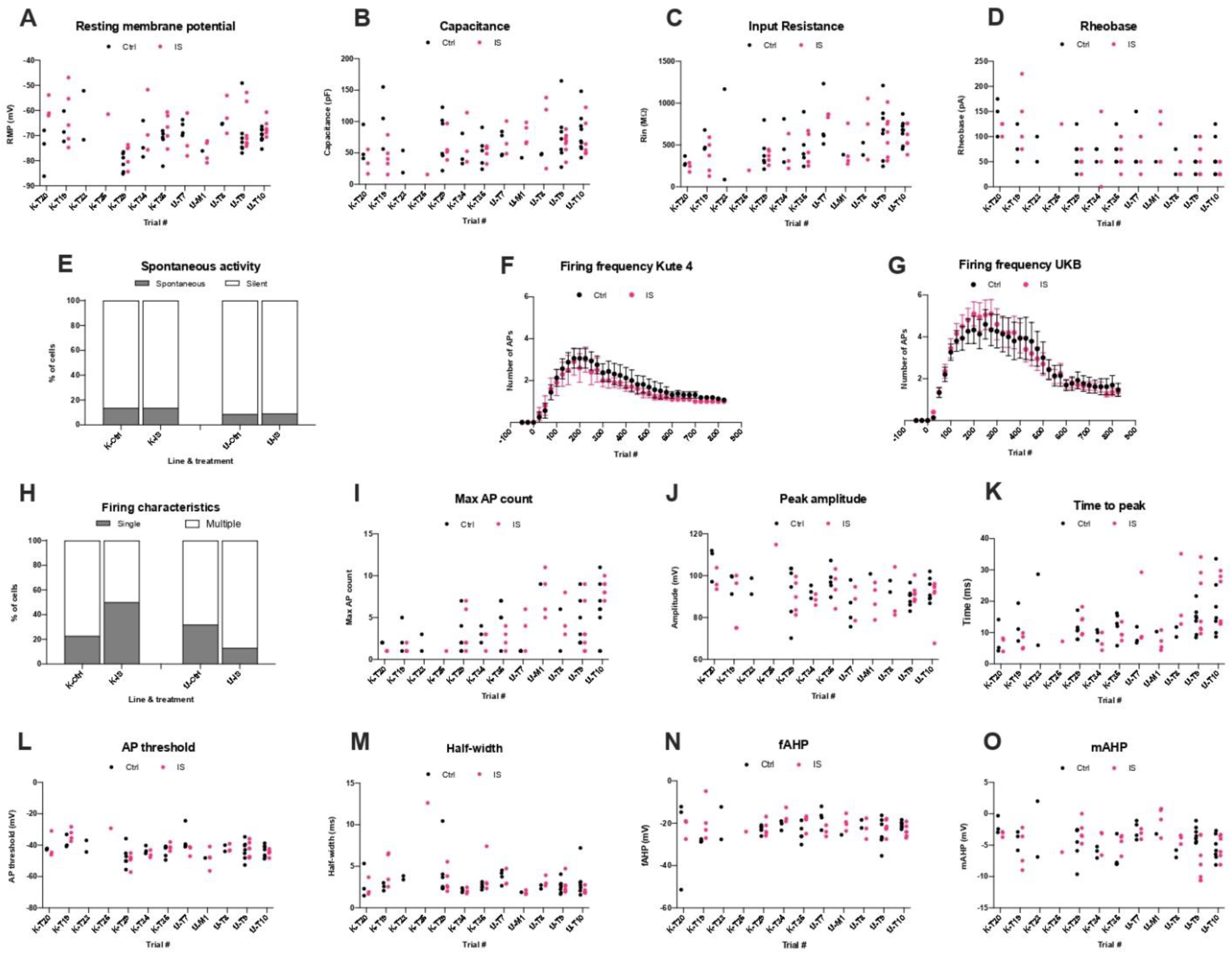
Electrophysiological characterisation of iPSC-derived sensory neurons in control and inflammatory conditions grouped per differentiation batch. The prefix “K” before the number stands for Kute4 line, “U” stands for UKB line. Black dots represent Ctrl, pink dots IS-treated cells. (A) Resting membrane potential, (B) capacitance, (C) Input Resistance, (D) Rheobase, (E) spontaneous activity, Firing frequency upon current injection for (F) Kute 4 and for (G) UKB, (H) firing characteristics, (I) number of maximal action potentials, (J) peak amplitude, (K) time to peak, (L) action potential threshold, (M) half-width, (N) fast afterhyperpolarisation (fAHP), (O) and medium afterhyperpolarisation (mAHP).

**Supplementary Figure 3.**
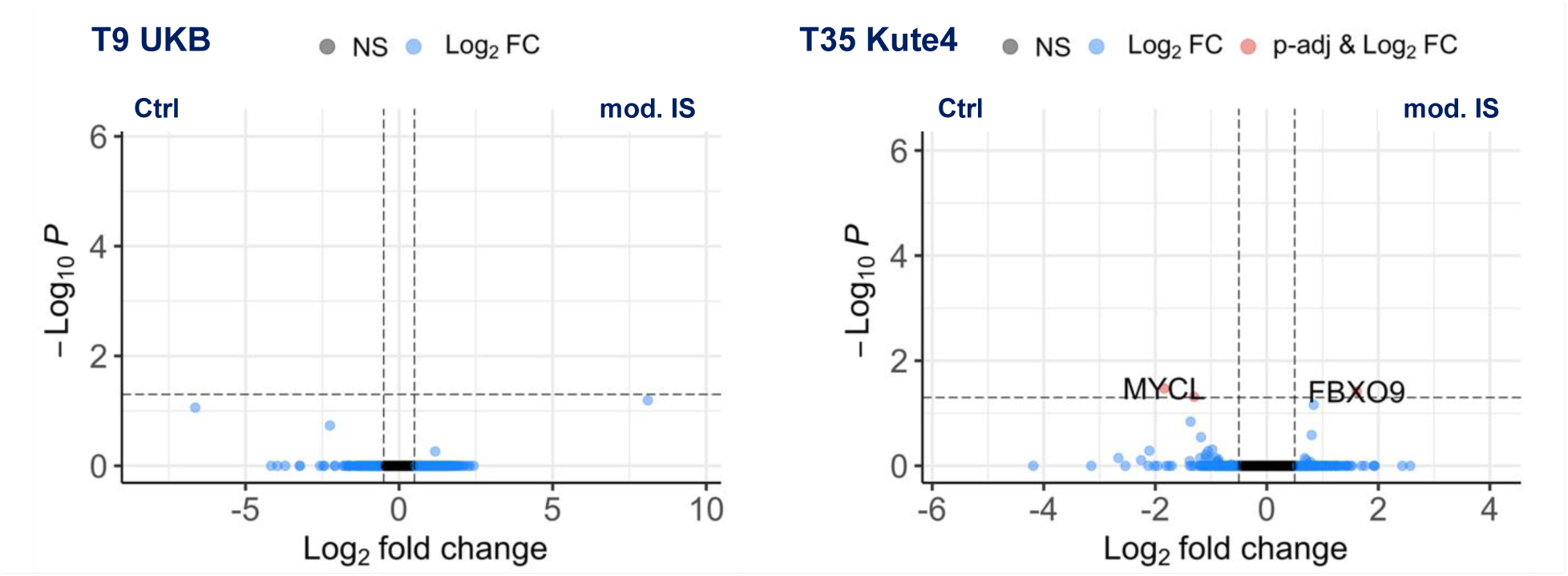
Volcano plots displaying differentially expressed genes for T9-UKB and T35-Kute4, respectively. padj cut-off: <0.05, Log_2_FC cut-off: > 0.5 or< -0.5.

## Footnotes

1 Since data are generated a few neurons at a time, with a series of experiments carried out over weeks or months, it is standard practice to plot some of the data ‘as you go along’, at which point one might decide to add more cells or to stop. This may seem intuitive, but statistically, this is a textbook example of sequential analysis, which requires controlling of Type 1 error rates in order to avoid unintentional p-hacking [22].

